# Missing Value Imputation using XGboost for Label-Free Mass Spectrometry-Based Proteomics Data

**DOI:** 10.1101/2021.04.08.438945

**Authors:** Jian Song, Changbin Yu

## Abstract

The label-free mass spectrometry-based proteomics data inevitably suffer from the problem of missing values. The existence of missing values prevents the downstream analyses which need a complete data matrix. Our motivation is to introduce the state-of-art machine learning algorithm XGboost to realize a method of imputation which can improve the accuracy of imputation. But in practical, XGboost has many parameters need to be tuned to deliver on its potential high performance. Although cross validation may find the best parameters, it is much time-consuming. Alternatively, we empirically determined the parameters to two kinds of base learners of XGboost. To explore the robustness and performance of XGboost based imputation with predetermined parameters, we conducted tests on three benchmark datasets. As a comparative, six common imputation methods were also experimented in terms of normalized root mean squared error and Pearson correlation coefficient. The comparative experimental results indicated that the XGboost based imputation method using the linear base learner is competitive to or out-performs its competitors, including the random forest based imputation, by achieving smaller imputation errors and better structure preservation under the empirical parameters for the three benchmark datasets.

## 1 Introduction

The label-free mass spectrometry-based (MS) quantitative method has become a major means of proteomics. But the proteome data matrix acquired by the label-free MS method may suffer from missing, accounting for biochemical and analytical to bioinformatics mechanisms. The missing rate frequently ranges between 10% to 50% [1]. Unfortunately, the downstream analyses of proteomics data such as clustering and supervised learning need complete data without missing values. Of course, one solution to the missing data problem is to repeat the experiment. But this strategy is expensive and time-consuming. Another option of ignoring missing values would dramatically reduce the size and completeness of the data and limit the researchers’ ability to infer information from the data. Thus, one of the major challenges of proteomics data studies is to impute this missing data appropriately.

## 2 Related works

Over the past decade, a variety of imputation algorithms have been developed and subsequently discussed in the literature [2]. For the sake of illustration, we assume ***O***_*m×n*_ to be a *m* × *n*-dimensional proteomics data matrix. Each row of ***O***_*m×n*_ is one observation represented by ***o***_1*×n*_, each column of ***O***_*m×n*_ means one protein variable denoted by ***υ***_*n×*1_, and ***υ***_*i,j*_ means the i-th observation in the j-th variable. Firstly, [3] published the first two missing value imputation algorithms based on the k-nearest neighbours (kNNImpute) and singular value decomposition (SVDImpute). The kNNImpute searches the most similarity observations and calculates the weighted sum as the imputation value. Mathematically,

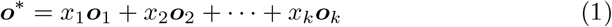

where ***o***^*\**^ is the targeted observation needed to be imputed, *k* (1 ≤ *k* ≤ *n*) is the selected observations’ numbers that mostly similarity to ***o***^*\**^. In most cases the *k* is the hyper-parameter needed to be determined by experiments. *x*_*i*_, *i* ∈ [1, *k*] is the corresponding coefficient between the selected observation with the target observation. The weight *x*_*i*_ is calculated as follow equation:

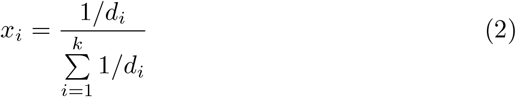

where *d*_*i*_ = ∥***o***_*i*_ − ***o***^*^ ∥_2_ is the Euclidean distance between ***o***_*i*_ and ***o***^*\**^. The SVDImpute first imputes the missing values with mean of variables to obtain a complete matrices, then repetitively decomposes the matrices and ignores the small singular value term until the total change in the matrix falls below the determined threshold. In math,

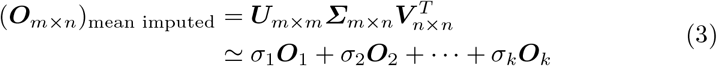

where ***U***, ***V*** is orthogonal matrix, and ***Σ*** is the diagonal matrix of singular values. Selecting the first *k* maximum singular values (*σ*_1_ ≥ *σ*_2_ ≥ … ≥ *σ*_*k*_) and calculating the sum can restore the raw matrix approximately.

[4] proposed a new missing value estimation method based on Bayesian principal component analysis (BPCAImpute). BPCAImpute consists of three elementary processes: (1) principal component regression, (2) Bayesian estimation, and (3) an expectation-maximization repetitive algorithm. BPCAImpute is a non-parameter method and can provide accurate and convenient estimation for missing values. Further, [5] and [6] introduced the least squares regression to imputation (LSImpute and LLSImpute). To simplify the description, we assume only the first element, i.e. *o*_1,1_ is missing, devoted by *η*. Therefore, the ***O*** can be divided into four parts:

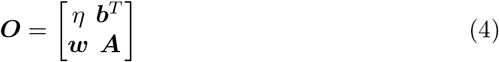

where the ***w*** and ***b***^*T*^ are the first column and row removed the *o*_1,1_ respectively and ***A*** is the remaining matrix removed the first row and column. Constructing the least squares regression model as

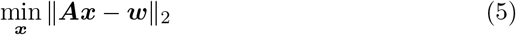

This is a well-know regression problem, and the coefficient vector ***x*** can be worked out by

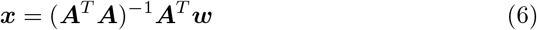

Then, the missing value *η* is estimated as

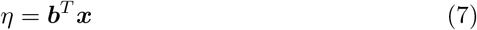

Same with this linear regression way, [7] and [8] trained the linear regression model equation (5) with regularization terms:

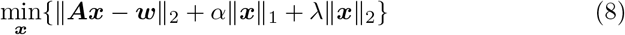

where *α* is the *L*_1_ regularized term coefficient, and *λ* is the *L*_2_ regularized term coefficient. [8] found that only reserving the *λ* can have better imputation result on test datasets. In this case, the *η* is given in

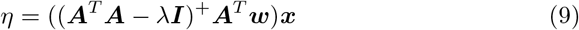

where the ***I*** is an identity matrix, and ()^+^ means to calculate the pseudo-inverse. After imputing the *o*_1,1_, other columns within missing values are estimated in the same process.

At present, ensemble learning categorized into bagging and boosting has shown its advantages in various fields for the application of classification and regression problem. Random forest is the typical of bagging method. [9] employed the random forest to model the ***A*** and ***w***. Based the trained model, the *η* can be predicted by ***b*** (RFImpute). According to [10] report, the random forest based imputation can achieve the minimum imputation error compared to other imputation methods. For the boosting method, typically gradient boosting machine (GBM) was firstly presented by [11]. and evolved to XGBoost [12]. In brief, the XGboost predicts the result with the sum of tree ensemble models:

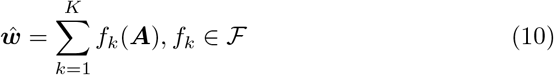

here *K* is the regression tree numbers and 𝒮 is the space of regression tree, *f*_*k*_(***A***) is the result of the *k*th tree. Then, we minimize the following regularized objective

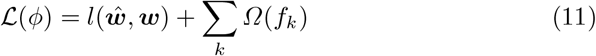

where the *l* denotes a differentiable convex loss function, *Ω*(*f*) is a regularization term that can penalize the model complexity. However, the tree ensemble model contains functions as parameters is unlikely to be optimized in Euclidean space when adapting traditional optimization methods. Thus, the model is trained as an effective manner that we can add *f*_*t*_ in the loss function. In this case, 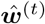 will become the prediction of the *t*th iteration. Hence, the regularized objective can be approximately described to the following formula of iteration *t*

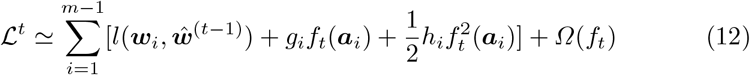

where ***a***_*i*_ is one row of ***A***, *f*_*t*_(***a***_*i*_) is the increment. 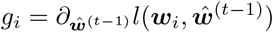 and 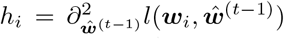 are the first and second order gradient statistics in the loss function. The model in XGboost is trained till stop criterion such as early stopping rounds or the number of boosting iterations *t* is satisfied. Besides the regression tree can be the base learner, the XGboost is also using the linear regression as a choice of base learner.

In this study, we modelled the ***A*** and ***w*** using XGboost, the gradient boost algorithm to improve the imputation accuracy for label-free mass spectrometry-based proteomics data. Meanwhile, we compared our method with meanImpute (imputation using constant mean value), kNNImpute, SVDImpute, BPCAImpute, LLSImpute and RFImpute. Comparisons were performed on three benchmark datasets coming from different papers and using different proportions of missing values.

## 3 Approach

### 3.1 Methods

The proteomics data are displayed as a matrix with rows and columns representing observations/samples and variables/proteins, respectively. Throughout the paper, we will use ***O***_*m×n*_ to denote the data matrix with m observations and n variables. First, let’s look at the case that only one column has the missing values. For instance, the first column is including missing values. Then we can divide the matrix into four parts as Fig.1 shows:

**Fig. 1.**
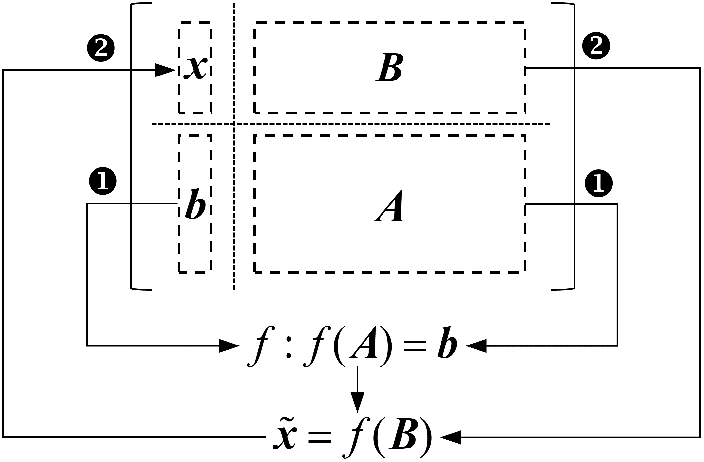
The illustration of data matrix division when only the first column need to be imputed.

1. the missing values vector of first column denoted as ***x***, and this is what we want to predict
2. the variables other than the first column with the same row index as ***x*** denoted by ***B***
3. the vector of first column excluding ***x*** denoted as ***b***
4. the remaining of matrix denoted by ***A***

After the definition of the matrix, the missing values ***x*** can be estimated through two phases: train and prediction. First, we use the matrix ***A*** and vector ***b*** to train the XGboost model, then, we feed the trained model with new data ***B*** to obtain the final estimated value.

For the case of missing values within multiple columns, we first need to make an initial guess for the missing values in ***O*** using mean imputation or another imputation method. Then, sorting the variables ***υ***_*i*_, *i* = 1, …, *n* according to the amount of missing values and starting imputation with the lowest amount missing values variable. For each variable ***υ***_*i*_, the missing values are imputed by the above process. After all the variables are imputed once, the imputation procedure is repeated until a stopping criterion is met. The pseudo Algorithm 1 exhibits a representation of the XGboost (xgbImpute) based imputation method inspired by [9].

The stopping criterion *γ* is met when the repetition times reaches a predetermined value or as soon as the difference between the newly imputed data matrix and the previous one increases for the first time. In that case, the *γ* is defined as

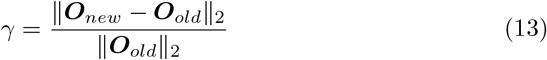

### 3.2 Evaluation Metrics

To evaluate the effectiveness of xgbImpute, normalized root mean squared error (NRMSE) [3] and Pearson correlation coefficient (PCC) [8] are used. The normalized root mean squared error measures the overall deviation of estimated values from their corresponding true value as defined below:

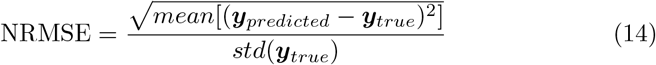

#### Algorithm 1 Impute missing values with XGboost

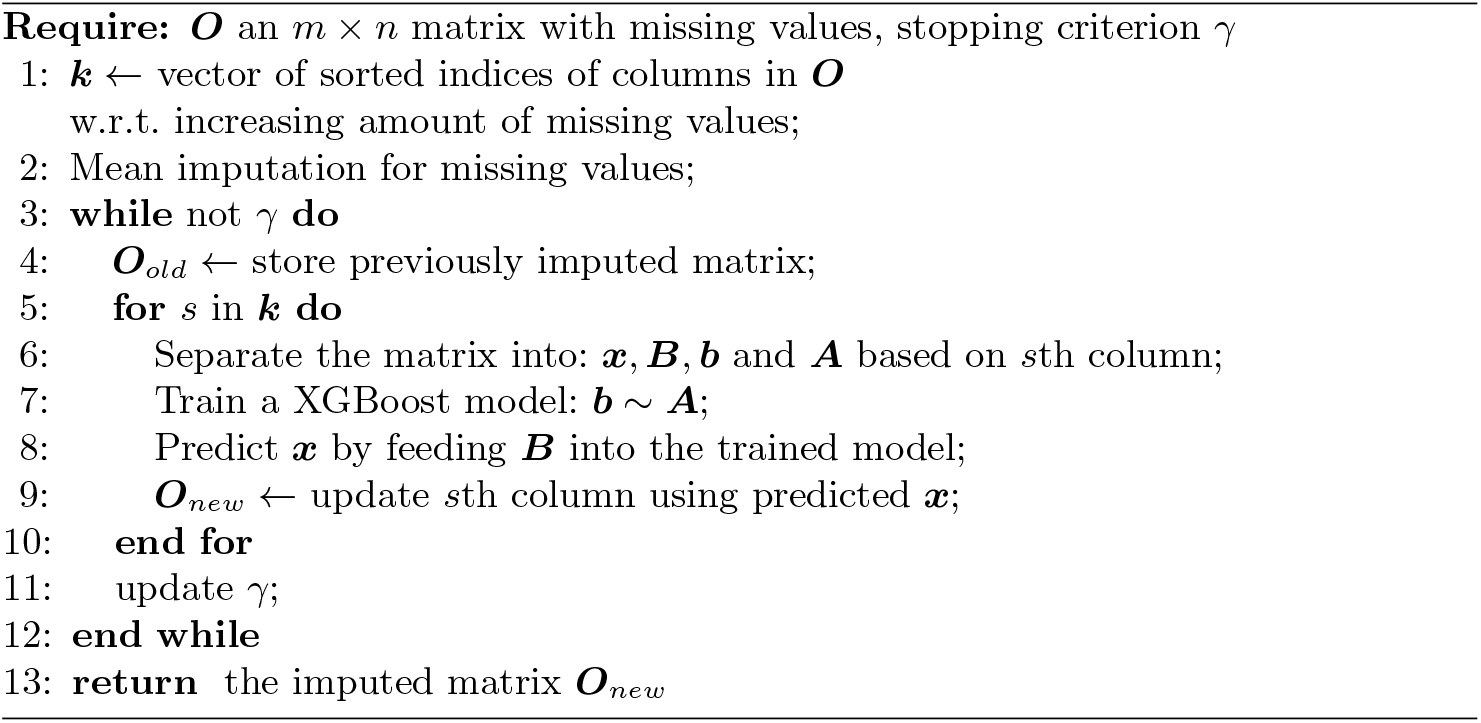

where ***y***_*true*_ is the vector of true value but is artificially missing, and ***y***_*predicted*_ is the vector of the predicted value correspond to ***y***_*true*_. NRMSE is more near to zero, the imputation is more accurate. Besides, Pearson correlation coefficient (PCC) is also able to quantify the consistence between the predicted and the raw data:

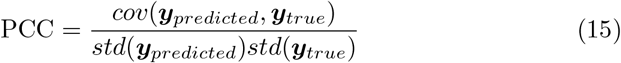

PCC is more near to 1, the imputation gets better accordance with actual data. These two metrics above reach a comprehensive comparison between xgbImpute and its competitors.

## 4 Experiments

### 4.1 Datasets

- Kinases expression of human colon and rectal cancer cell line (CRC): the authors used LC-MS/MS-based shotgun proteomics to measure the proteomes of 65 CRC lines and to quantify their expressed kinomes using Kinobeads ([13], [14]). This led to the identification of a total of 235 human kinases (median/cell line = 155, from Kinobeads experiments), see [15]. The missing rate of this dataset is about 34%.
- Proteome about the interstitial lung disease (ILD): This dataset has been previously described in detail by [16]. From each sample in 11 human lung tissue biopsies, the proteins were extracted with increasing stringency into four fractions by changing the detergent and buffer conditions. We select the result of first fraction and randomly sample 500 proteins without missing values.
- Ovarian cancer proteome dataset (Ovarian): The authors performed quantitative proteomics on 25 ovarian tumor samples. Adapting label-free proteomic workflow they identified and quantified more than 9,000 proteins from low-mg archival samples in single-run measurements in the MaxQuant environment. More details see [17]. Randomly we select 400 proteins to construct the complete data matrix.

For the ILD and Ovarian dataset, we randomly introduced missing values to the complete dataset with missing rates of 10% to 50% stepped with 10%. Meanwhile, we saved the true value to evaluate the imputation accuracy later. For the CRC datasets, accounting to the original missing rate, we introduce missing values with missing rates from 35% to 50% stepped with 5%. Due to the missing mechanism is random, we performed 10 times imputation experiments for every missing rates.

### 4.2 Parameters

We compared xgbImpute with meanImpute, kNNImpute, SVDImpute, BPCAImpute, LLSImpute and RFImpute. The detail information of these methods was shown in Table 1. The parameters of k nearest neighbors in KNNimpute, the number of components in SVDImpute and the number of similar variables in LLSImpute are hyper parameters and we tried all possible parameters in [1, *m*− 1] under the first missing rate of each dataset and selected the parameter in the best performance with the metrics of NRMSE.

**Table 1.**
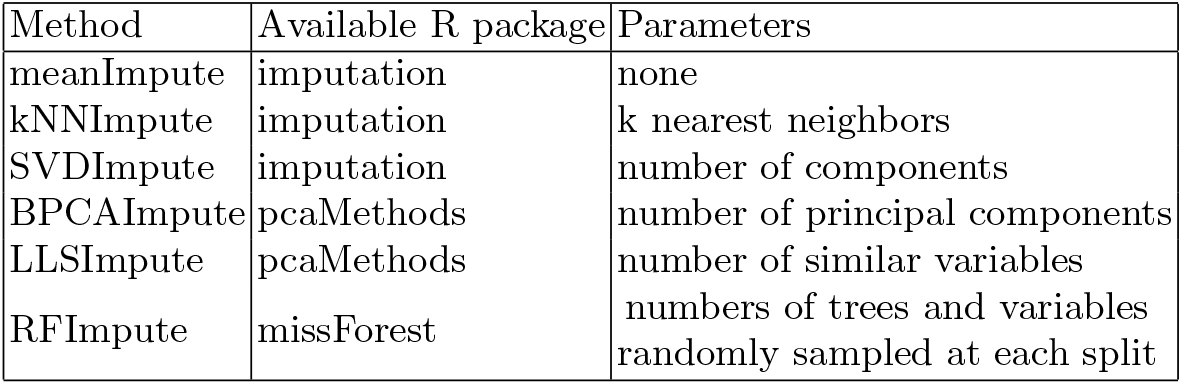
Competition Methods Description

The parameter of BPCAImpute is the number of principal components which was set to the numbers of rows subtracted one as recommended in [4]. The parameters of RFImpute include the numbers of trees and variables randomly sampled at each split for both of which we used the default values.

The parameters of xgbImpute are inherited from XGboost. When the base learner is regression tree, i.e. `booster`= ‘gbtree’, the parameters mainly including:

- `nrounds` max number of boosting iterations.
- `eta` control the learning rate. It is used to prevent overfitting by making the boosting process more conservative.
- `gamma` minimum loss reduction required to make a further partition on a leaf node of the tree.
- `max depth` maximum depth of a tree.
- `min child weight` minimum sum of instance weight (hessian) needed in a child.
- `subsample` subsample ratio of the training instance.
- `colsample bytree` subsample ratio of columns when constructing each tree.

When the base learner of XGboost is linear model, i.e. `booster` = ‘gblinear’, the parameters mainly including:

- `nrounds` max number of boosting iterations.
- `lambda` L2 regularization term on weights.
- `lambda bias` L2 regularization term on bias.
- `alpha` L1 regularization term on weights.

In order to optimize parameters comprehensively, we consulted some references ([12], [18], [19]) and determined by grid search with cross validation. Ultimately, the results we used are : `nrounds` = 100, `eta` = 0.15, `gamma` = 0, `max depth` = 6, `min child weight` = 1, `subsample` = 0.8, `colsample bytree` = 0.8 and parameters for the ‘gblinear’ booster are: `nrounds` = 100, `lambda` = 10, `lambda bias` = 0 and `alpha` = 0.

## 5 Results

### 5.1 normalized root mean squared error

In this study, for each dataset, we conducted experiments ten times for each imputation method and each missing rate and calculated the average normalized root mean squared error(the xgbImpute including two methods based on the base learner: gbtreeImpute and gblinearImpute). As shown in Fig. 2, we can observe that NRMSE increases with the increasing of missing rates for all imputation methods approximately. This is reasonable because a larger missing rate comes with the loss of more information. Besides the average NRMSE, we also performed a unpaired Wilcoxon test of the NRMSE of the compared methods versus the NRMSE of gblinearImpute which have the best performance in most datasets at each missing rate.

**Fig. 2.**
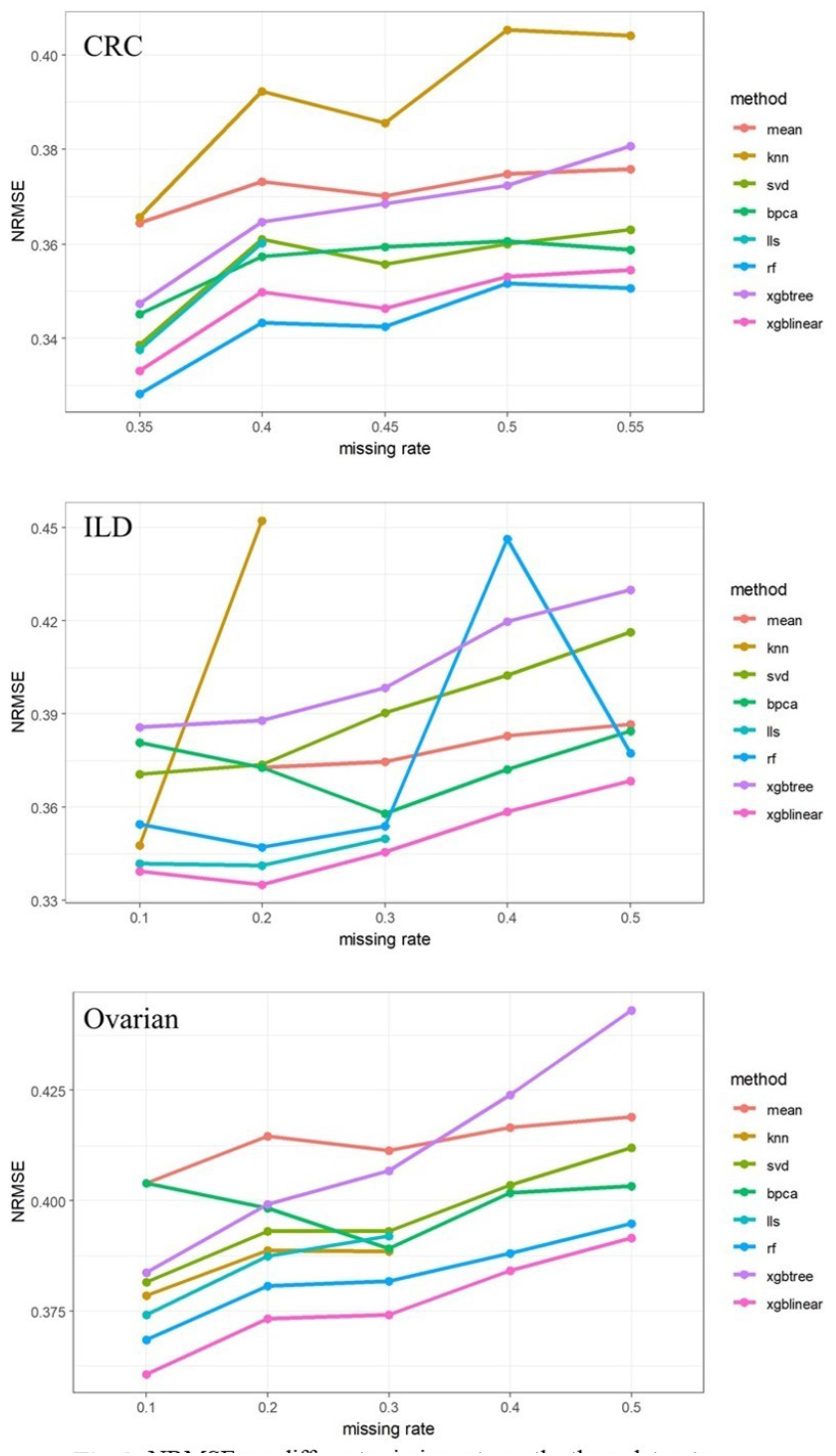
NRMSE vs. different missing rate on the three datasets.

Combining the average NRMSE result and significance result of CRC dataset in Table 2, we can see that the RFImpute have the minimum average NRMSE at each missing rate. Meanwhich, none of the significance between RFImpute and gblinearImpute results are significant as Table 2 shows. That is to say, the performance of RFImpute and gblinearImpute for the CRC dataset are close to each other. In addition, the gbtreeImpute have the moderate performance compared to other methods. In the case of 45%, 50%, 55% missing rates, the LLSImpute algorithm produced huge NRMSE. To better display of other valid value, we don’t show these outliers.

**Table 2.**
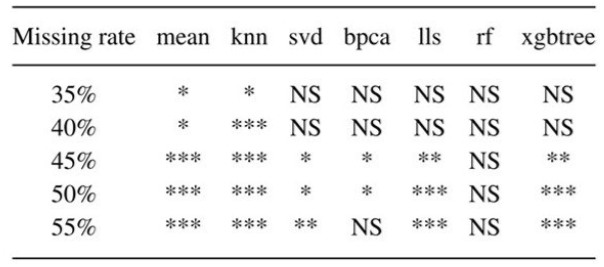
Results of Significance Test on CRC Dataset

For the ILD dataset, result shown in Fig. 2 (ILD) and Table 3, we can observe that the gblinearImpute have the best performance at each missing rate. Oppositely, the gbtreeImpute is not work better than other methods. Considering the significance result, we found kNNImpute, LLSImpute and RFImpute have no significance at missing rate 10% to 30% compared to gblinearImpute. But when the missing rate is increasing to 40% and 50%, the gblinearImpute is better than all other competitors significantly. Also, the kNNImpute and LLSImpute produce outliers of imputation result, and the RFImpute have a enormous NRMSE at missing rate 40%.

**Table 3.**
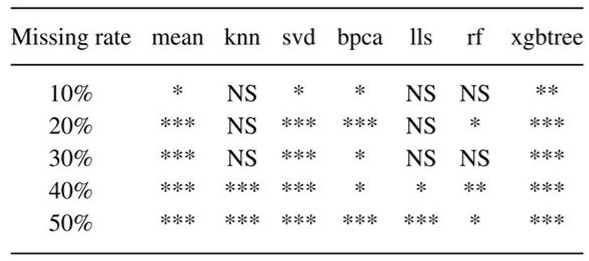
Results of Significance Test on ILD Dataset

For the Ovarian dataset, result shown in Fig.2 Ovarian and Table 4, we can see that the gblinearImpute is better than other methods significantly at each missing rate except the RFImpute at missing rate of 10% and 40%. Meanwhile, the gbtreeImpute has poor performance at missing rate 40% to 50% compared to other methods. In the case of 40%, 55% missing rates, the knnImpute algorithm produce huge NRMSE compared to other methods which are not shown.

**Table 4.**
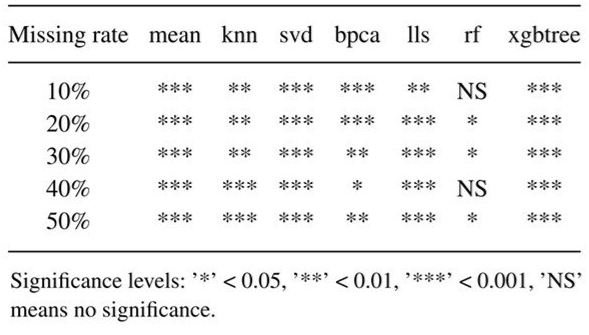
Results of Significance Test on Ovarian Dataset

Looking at the metric of NRMSE, the gblinearImpute with specify parameters is mostly better than other competitors at each missing rate. Considering the robust, the gblinearImpute is also consistently outperforms its competitors. For the specify parameters gbtreeImpute, it doesn’t show its advantages.

### 5.2 Pearson correlation coefficient

To show the effectiveness of different imputation methods in preserving the original data structure, we also calculated the PCC based on the true value and the predicted value. Fig.3 presents corresponding experimental results for different methods about the PCC values. Combining Fig.3 with Fig.2, we can observe that the average PCC trend is opposite to the average NRMSE trend, i.e. PCC decreases with the increasing of missing rates for all imputation methods approximately. In particular, we found that only in the case of 35% missing rate of CRC dataset, the performance of meanImpute and kNNImpute are inconsistent under the metrics of NRMSE and PCC. That is to say, in the view of NRMSE, the meanImpute is superior, but in the view of PCC, it is inferior. Except this case, we don’t find that the NRMSE is outperformance but the PCC is under-performance. Thus, looking at the metric of PCC, the conclusion is basically the same with the metric of NRMSE, i.e. for CRC dataset, the gblinearImpute and rfImpute is outperformance, for ILD and Ovarian dataset, the gblinearImpute is more accuracy and robust than other competitors.

**Fig 3.**
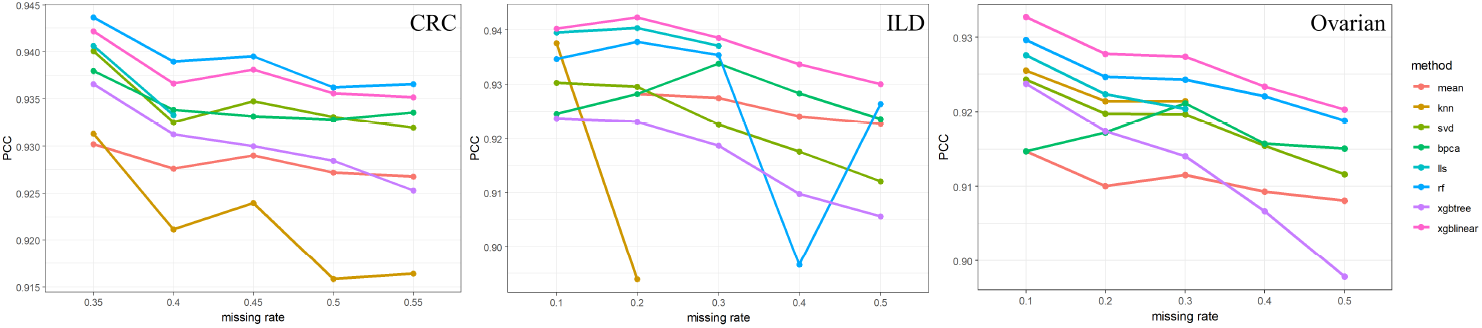
Pearson correlation coefficient vs. different missing rate on the three datasets.

## 6 Conclusion

Missing value imputation is a challenging for proteomics research. With the development of many kinds of imputation methods, it is more and more difficult to improve the accuracy and robustness of imputation method. In this paper, we applied the state-of-art machine learning algorithm XGboost to realize the imputation of label-free mass-spectrometry based proteomics data. Theoretically, when training the XGboost model every time for each missing column we can use cross validation to get the best parameters, but this will lead to much time-consuming. Alternatively, we empirically determined the parameters for the two kind base learners of XGboost. Our preliminary experimental results show on three real label-free mass-spectrometry based proteomics datasets that the allocated parameters of XGboost with linear base learner for imputation is competitive to or outperforms six established imputation methods like the random forest based imputation.

For subsequent analysis, we plan to analysis every sample predict error other than the whole error to find the cause of relative big mistake such as the distribution of missing values. Furthermore, we will explore the XGboost based imputation can be applied to other fields that also suffer from the problem of missing values such as DNA microarray or metabolic data.

